# EnzySeek: Efficient Exploration of Enzyme Reaction Pathways Using AI Agents

**DOI:** 10.64898/2026.03.02.708939

**Authors:** Xu Kang, Tao Yu, Kangwei Xu, Chenxu Liu, Ruibo Wu

**Affiliations:** School of Pharmaceutical Sciences, Sun Yat-sen University, Guangzhou 510006, P.R. China

## Abstract

With the rapid development of Large Language Models (LLMs) and Agent technologies, AI can assist in solving a variety of real-world problems across multiple domains, such as autonomous driving, drug discovery, and materials design. In this work, we present EnzySeek, an enzyme catalysis AI agent designed to assist researchers in enzyme catalysis simulations. First, we constructed a domain-specific knowledge base by curating thousands of papers related to enzyme catalysis. Second, we customized Model Context Protocol (MCP) interfaces for each step of the enzyme catalysis simulation workflow, enabling these functions to be invoked by LLMs. Finally, we configured an agent capable of simultaneously referencing past empirical studies on enzyme catalysis, autonomously executing tool calls, and analyzing as well as presenting the results.

EnzySeek’s capabilities cover multiple aspects, including protein structure prediction, molecular docking, system preparation and parameterization, molecular dynamics (MD) simulations, and QM/MM calculations. The conclusions drawn by EnzySeek are primarily based on the results of QM/MM calculations. We employed the semi-empirical quantum mechanical method GFN2-xTB to calculate the QM region of the system. Benchmark results indicate that the GFN2-xTB method can achieve high efficiency while maintaining accuracy.

The EnzySeek agent is designed to continuously learn from newly published literature and past computational tasks. During its operation, every AI decision is manually verified and scored by human experts. This human-in-the-loop validation provides the AI with sufficient case-based support, ultimately contributing to the full automation of enzyme catalysis computations. All data generated during the simulations are compiled into a dataset, which is used to establish evaluation criteria specific to enzyme catalysis computational results.

## Background

Enzymes[1, 2] are natural catalysts capable of catalyzing the synthesis of numerous compounds that are difficult to produce via traditional chemical methods. However, natural enzymes often face challenges in industrial or clinical applications, such as low catalytic efficiency, poor substrate specificity, and numerous side reactions. To overcome these limitations, enzymes can be engineered through approaches like rational design[3] or directed evolution[4], thereby significantly improving their catalytic efficiency and reducing byproduct formation. Nevertheless, enzyme engineering is typically associated with high experimental costs and long development cycles, severely hindering the research and development process.

Computational simulation technologies offer an accelerated pathway to address this issue. Among these, the combination of molecular dynamics (MD)[5, 6] simulations and quantum mechanics/molecular mechanics (QM/MM)[7, 8] methods has become a mainstream approach for investigating enzyme catalytic mechanisms due to its high accuracy and universality. However, in practical applications, MD and QM/MM simulations rely heavily on specialized knowledge in computational chemistry and expert experience, which greatly limits the widespread adoption of these techniques. Furthermore, the quantum chemical methods employed in QM/MM require meticulous calculations of electronic structures, resulting in exceedingly long computation times. Consequently, the simulation cycle for a single enzymatic reaction often spans 1 to 6 months. Therefore, the cumbersome operational and result-analysis workflows, combined with lengthy electronic structure calculations, collectively bottleneck the efficiency of enzyme catalysis simulations.

In this work, we replaced the high-precision quantum chemical methods in the conventional QM/MM workflow with the semi-empirical quantum mechanical method GFN2-xTB[8], and conducted detailed benchmarking on three distinct systems. The results demonstrate that the semi-empirical quantum mechanical method can qualitatively reproduce the computational results of high-precision methods while reducing the computation time by three orders of magnitude, thereby significantly shortening the cycle of a single QM/MM calculation. Consequently, under the QM(GFN2-xTB)/MM framework, the same computational power can simultaneously support thousands of times more computational tasks. At this scale, the time consumed by manual operations and result analysis becomes particularly prominent. By leveraging Agent technology[9], we encoded all common computational and result-analysis workflows into functions callable by LLMs[10], unifying all function interfaces via the Model Context Protocol (MCP)[11, 12]. We also utilized a system engine to comprehend computational tasks, execute calculations, and analyze results, significantly reducing the human burden associated with submitting computational tasks and processing outcomes.

The EnzySeek agent consists of three modules: a Knowledge Base, a Skill Base, and a Dataset. The Knowledge Base aggregates relevant literature in the field of enzyme catalysis, allowing the agent to reference similar cases when executing specific tasks. The Skill Base pre-provisions functional routines for the entire enzyme catalysis workflow, empowering the agent to execute real computational tasks, acquire data, and analyze it. The Dataset serves as the agent’s memory, preserving historical operational workflows and structural data to construct criteria for evaluating the rationality of enzyme catalysis computations. In this paper, we demonstrate EnzySeek’s capability to solve practical problems through two previously published cases.

Our main contributions are as follows:

- Evaluated the applicability of GFN2-xTB in QM/MM calculations of enzymatic reactions: We calculated the energies and structures of enzyme-ligand complexes across three published systems. We found that semi-empirical (SE) QM/MM can qualitatively reproduce the results of *ab initio* QM/MM while reducing the time consumption by three orders of magnitude.
- Proposed an implementation framework for Agent technology in specialized domains: In our study, we successfully built a domain-specific agent by first gathering existing knowledge (primarily research papers), then encoding domain-specific operational workflows into LLM-callable functions, and finally continuously optimizing the model’s decision-making through human-AI collaboration.
- Developed the enzyme catalysis Agent, EnzySeek: The enzyme catalysis agent we developed incorporates a comprehensive collection of literature in the field of enzyme catalysis, extracting knowledge and computational results from it to assist in decision-making. It significantly reduces manual labor costs by executing high-throughput computational tasks and summarizing the analytical results.

## Results

### Architecture and Workflow

The platform is primarily divided into a backend server responsible for data storage, task scheduling, and decision-making, and a client-side interface dedicated to result visualization and human-computer interaction.

The backend server (Figure 1 left) consists of three core modules: a Knowledge Base, a Skill Base, and a Dataset. The Knowledge Base extracts enzyme catalysis-related information from domain-specific literature. Each piece of knowledge is indexed based on features such as the enzyme, reaction type, and computational method, facilitating the retrieval of computational details and key conclusions. The Skill Base encompasses various modules covering the entire enzyme catalysis simulation workflow, including protein modeling, structural optimization, molecular docking, molecular dynamics (MD), and potential energy surface (PES) scans. These tools are unified under the Model Context Protocol (MCP) for standardized input and output, enabling direct invocation by Large Language Models (LLMs). The Dataset archives all accessible computational results within the domain, as well as the historical task execution logs of the system, which are used to construct a comprehensive evaluation framework for simulation outcomes.

**Figure 1.**
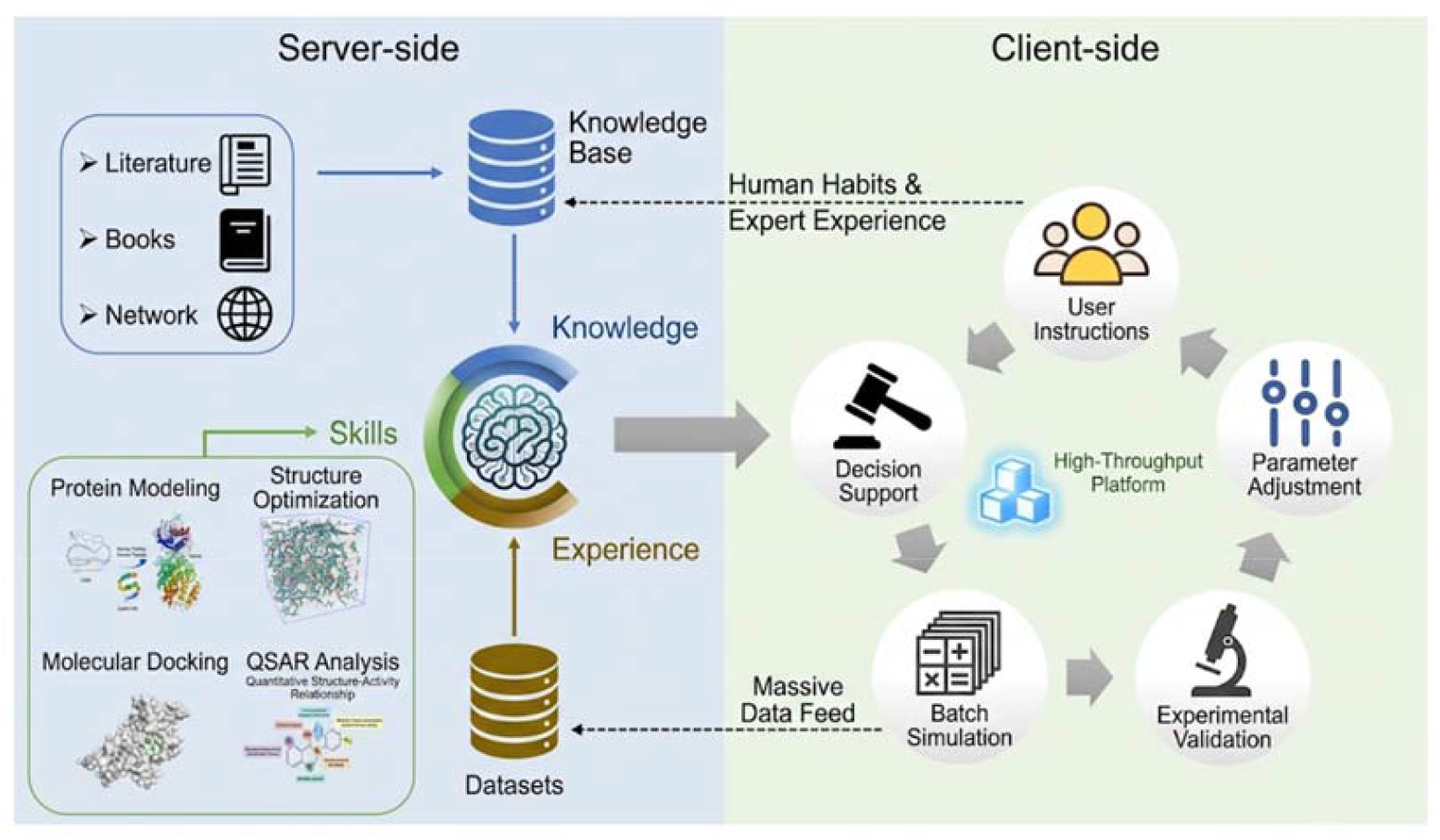
Overall architecture of the EnzySeek platform. The server-side (left) encompasses the agent module, which is responsible for knowledge storage and tool invocation. The client-side (right) features a Web UI dedicated to result visualization and human-agent interaction.

The client (Figure 1 right) operates via a Web UI for human-computer interaction. Human researchers can invoke the system’s tools to execute calculations and retrieve results using natural language. Furthermore, researchers can correct unreasonable AI-generated decisions and fine-tune various AI-recommended parameters based on their expert knowledge. For systems with existing experimental data, researchers can preemptively supply these results to provide heuristic guidance for the agent.

### Knowledge Base and Dataset

We curated thousands of research articles and their supporting materials related to enzyme catalysis simulations. Utilizing a multimodal LLM, we parsed the text, figures, and tables from these documents, compiling all experimental details and results into the Knowledge Base. We constructed exclusive indices for each knowledge entry based on enzyme type, ligand type, and reaction type. These indices are prioritized during retrieval when the agent processes similar tasks, ensuring the recall of highly relevant reference information. Additionally, all authentic operational logic and instructions provided by human researchers are stored in the Knowledge Base. These data assist the agent in learning human behavioral patterns and the implicit expert knowledge embedded within them.

For all completed computational tasks, human researchers evaluate the results. High-quality outcomes—such as the volume of the enzyme binding pocket, the active site of a specific system, and the dominant conformations of different substrates—are deposited into the Dataset. These data are further analyzed and abstracted into evaluation criteria for future computational results. Consequently, when the model encounters similar systems in the future, it can autonomously submit tasks and analyze results by referencing the computational methodologies in the Knowledge Base and the evaluation standards in the Dataset.

### Functions

We encoded every step of the enzyme catalysis workflow into distinct functions. By employing the MCP to standardize input and output specifications, the LLM can efficiently invoke each predefined function and obtain standardized computational results. In short, we defined LLM-callable interfaces for the following key functionalities:

- **Protein Structure Prediction:** Utilizes Chai-1[13] to predict 3D structures based on provided protein sequences. It also supports the autonomous upload of PDB files for subsequent workflows.
- **Molecular Docking:** Employs AutoDock[14] to perform protein-ligand docking, returning the top ten highest-scoring complexes. It also accepts pre-processed protein-ligand complex structures.
- **Solvation:** Uses the *tleap* module in AmberTools[15] for system solvation and force field parameterization.
- **QM/MM Calculations:** Utilizes AMBER[16] for the partitioning of the QM and MM regions. The QM region is calculated using the GFN2-xTB method. By default, the QM region includes the ligand and residues within 5 Å of the ligand. If metal ions are present in the pocket, residues within 3 Å of the metal ions are also included. The maximum number of atoms in the QM region is capped at 400; if this limit is exceeded, residues are sequentially removed based on their distance from the ligand, starting from the furthest.
- **Potential Energy Surface (PES) Scan:** Applies restraint keywords to constrain specific bond lengths, bond angles, or dihedral angles. Restrained optimizations are performed by relaxing the remaining degrees of freedom, and all converged energies are extracted as the final result.
- **QM/MM MD Simulations:** Performs MD simulations at the QM/MM level using pre-parameterized systems, with a default simulation time of 50 ps.
- **Result Analysis:** Utilizes custom scripts to parse structural data, energies, and dynamic trajectories from AMBER output files.

### Benchmark

The stability of the method was benchmarked using previously published systems from our group, including *Pc*TS1[17], EfTPS14[18], and VenA[19]. The evaluation metrics encompassed differences in calculated energy barriers and enthalpy changes for identical reaction pathways across different methods, as well as root-mean-square deviations (RMSD) of the optimized ligand conformations.

Table 1 presents the calculated energy barriers and enthalpy changes for representative reactions in the *Pc*TS1 and EfTPS14 systems. We recalculated the energies of the intermediate structures reported in the literature at the GFN2-xTB level and compared them with the published data. The results indicate that GFN2-xTB can qualitatively reproduce the computational results of high-precision quantum chemistry methods.

**Table 1.** Calculated energy barriers and energy changes for key reaction steps in the pcts1 and EFsTPS14 systems (Units: kcal/mol)

| **System** | **$\Delta E^\ddagger$ (M062X)** | **$\Delta E^\ddagger$**<br>**(GFN2-xTB)** | **$\Delta E$**<br>**(M062X)** | **$\Delta E$**<br>**(GFN2-xTB)** |
| --- | --- | --- | --- | --- |
| *PcTS1-1* | / | / | -10.0 | -13.0 |
| *PcTS1-2* | / | / | -8.0 | -20.0 |
| EfTPS14-1 | 14.0 | 8.4 | 7.5 | 2.2 |
| EfTPS14-2 | 1.4 | 0.5 | -22.4 | -24.4 |
| EfTPS14-3 | 12.0 | 7.0 | -1.9 | -2.4 |

In the structural comparison of the VenA enzyme-ligand complex, structural optimization was initiated from the key intermediates reported in the literature. Optimization was performed at the semi-empirical QM/MM level, and the RMSD between the converged geometry and the initial structure was calculated. We found that the average RMSD difference between the QM/MM optimized structures at the GFN2-xTB level and those at the M062X level was less than 1 Å (Figure 2). The corresponding energies also qualitatively reproduced the energy profiles of key intermediates reported in the literature (Table 2).

**Table 2.**
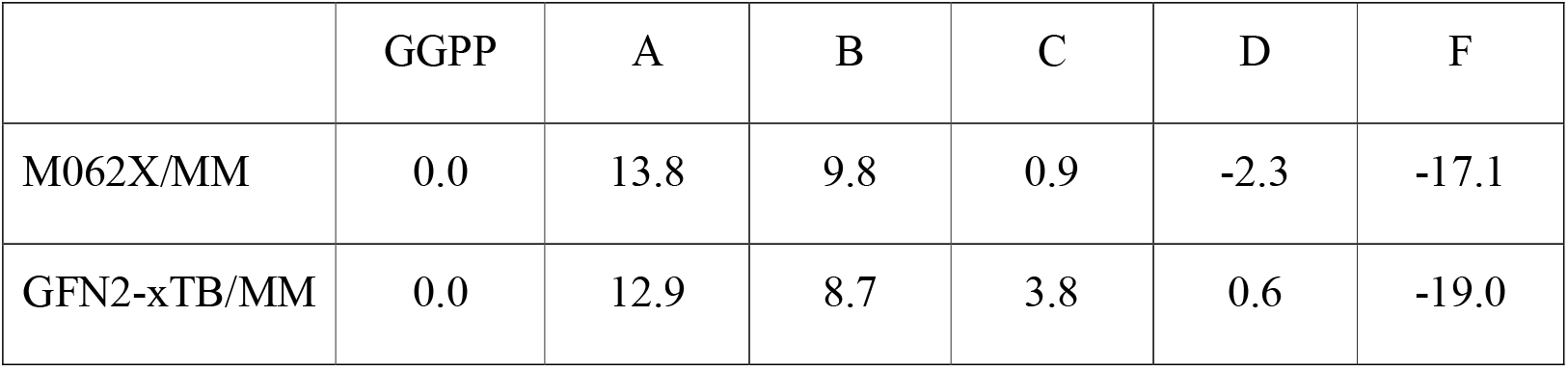
Calculated energies of different intermediate structures in the VenA system using two distinct QM/MM methods (units: kcal/mol).

**Figure 2.**
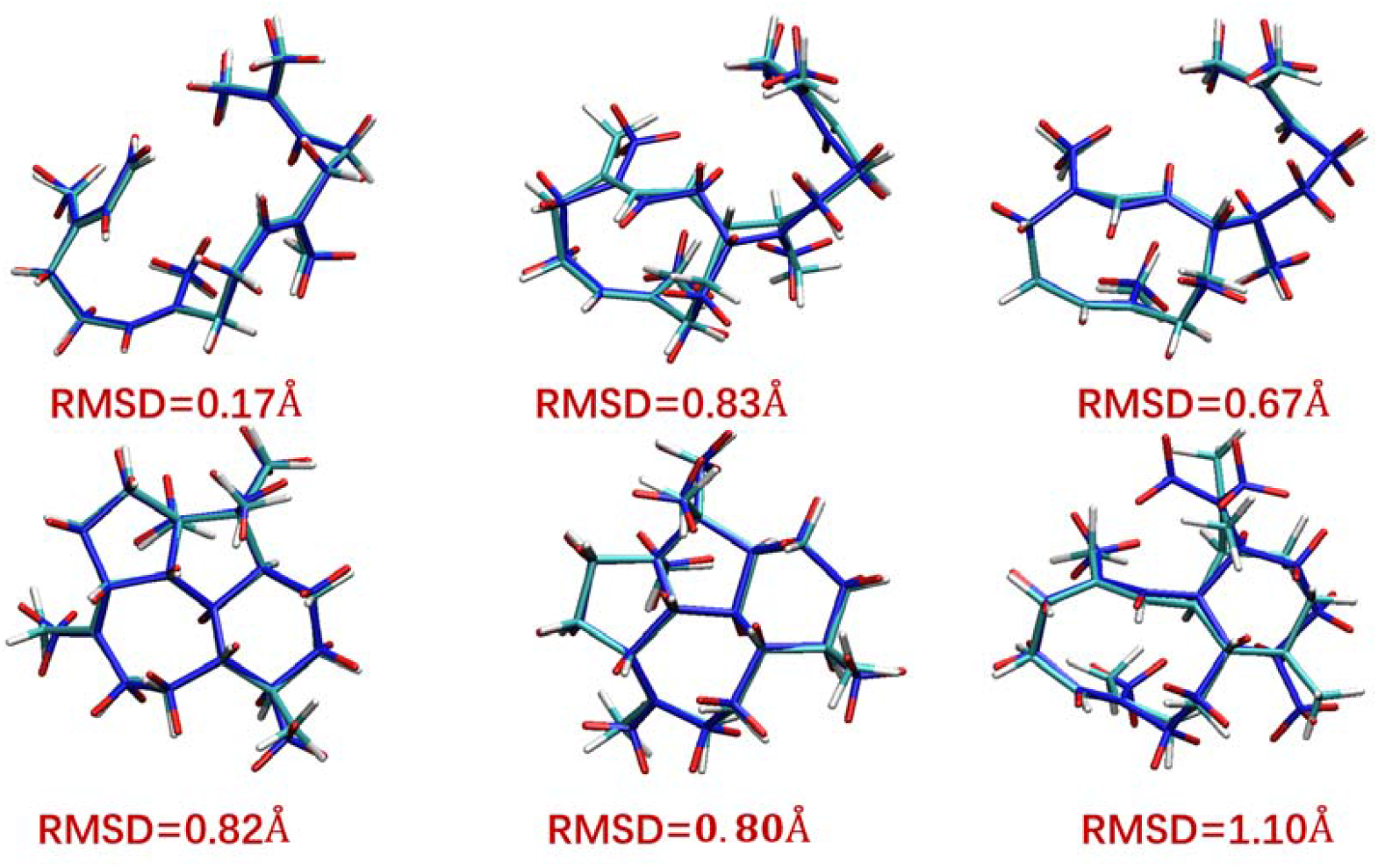
Comparison of the converged ligand conformations for different intermediates in the VenA system following QM/MM structural optimization using two distinct QM methods. Cyan structures represent optimizations performed at the M062X level, while blue structures correspond to those optimized using the GFN2-xTB method.

### Case Study

#### Case 1: Sampling Dominant Substrate Conformations in Synthases

This case investigates the sampling method for dominant substrate conformations in a synthase[20]. In this system, the synthetic route depends heavily on the initial binding mode of the substrate and the folding tendency of the ligand within the pocket.

Traditional MD methods struggle to accurately describe subtle conformational changes of the ligand within the enzyme pocket, whereas high-precision QM/MM methods are too computationally expensive to observe over sufficiently long timescales. Therefore, we employed the EnzySeek Agent to conduct QM/MM MD simulations to identify and summarize the dominant conformations of the ligand within the system.

1. First, EnzySeek invoked the molecular docking module to dock 4 potential key reaction intermediates into the pocket. For each intermediate, the top 10 results with the lowest energies were selected, yielding a total of 40 initial systems (Figure 3B).
2. Next, EnzySeek performed QM/MM MD simulations on these 40 initial frames at the GFN2-xTB level.
3. Finally, EnzySeek conducted conformational analysis on the final frames of all QM/MM MD trajectories, successfully simulating the dominant conformations of the ligand in the target system (Figure 3C).

**Figure 3.**
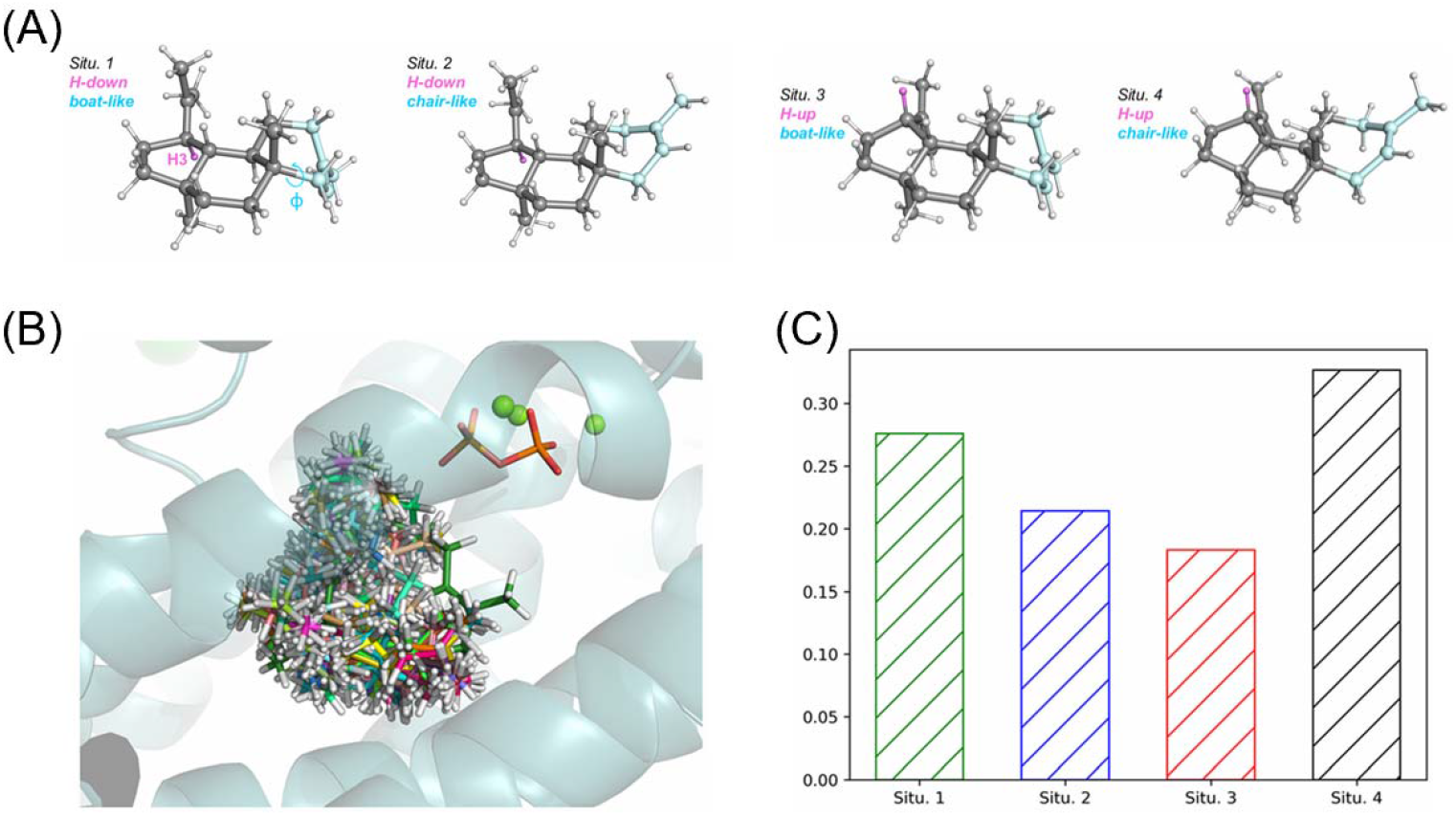
Key results of EnzySeek in the search for the dominant reaction pathway of Cyathane. (A) Four primary conformations of Cyathane found in nature. (B) Forty conformations obtained via molecular docking of the four structures, which serve as the initial frames for the subsequent QM/MM MD trajectories. (C) Bar chart illustrating the classification of the four conformations by EnzySeek following the completion of the QM/MM MD simulations.

#### Case 2: Investigating Catalytic Promiscuity in Isoflavone-4’-O-methyltransferase

This case addresses the catalytic promiscuity of Isoflavone-4’-O-methyltransferase[21]. This enzyme is a critical component in the biosynthetic pathway of the dihydroisoflavone compound medicarpin, responsible for catalyzing the 4’-hydroxyl methylation of 2,7,4^′^-trihydroxyisoflavanone. Experimental studies have revealed that this enzyme can also catalyze the methylation of another flavonoid substrate, liquiritigenin, at both the 4’- and 7-hydroxyl positions. To explore this catalytic promiscuity, we conducted simulations to elucidate the underlying reaction mechanisms.

The catalytic mechanism involves the active-site residue H270 abstracting a proton from the substrate’s hydroxyl group, while the hydroxyl oxygen simultaneously mounts a nucleophilic attack on the methyl donor SAM, yielding the methylated product. Additionally, residue E332 forms a hydrogen bond with H270, enhancing the basicity of the histidine. Thus, whether the substrate’s hydroxyl group can maintain an appropriate reaction distance and orientation relative to SAM and H270 determines the feasibility of the reaction (Figure 4A).

**Figure 4.**
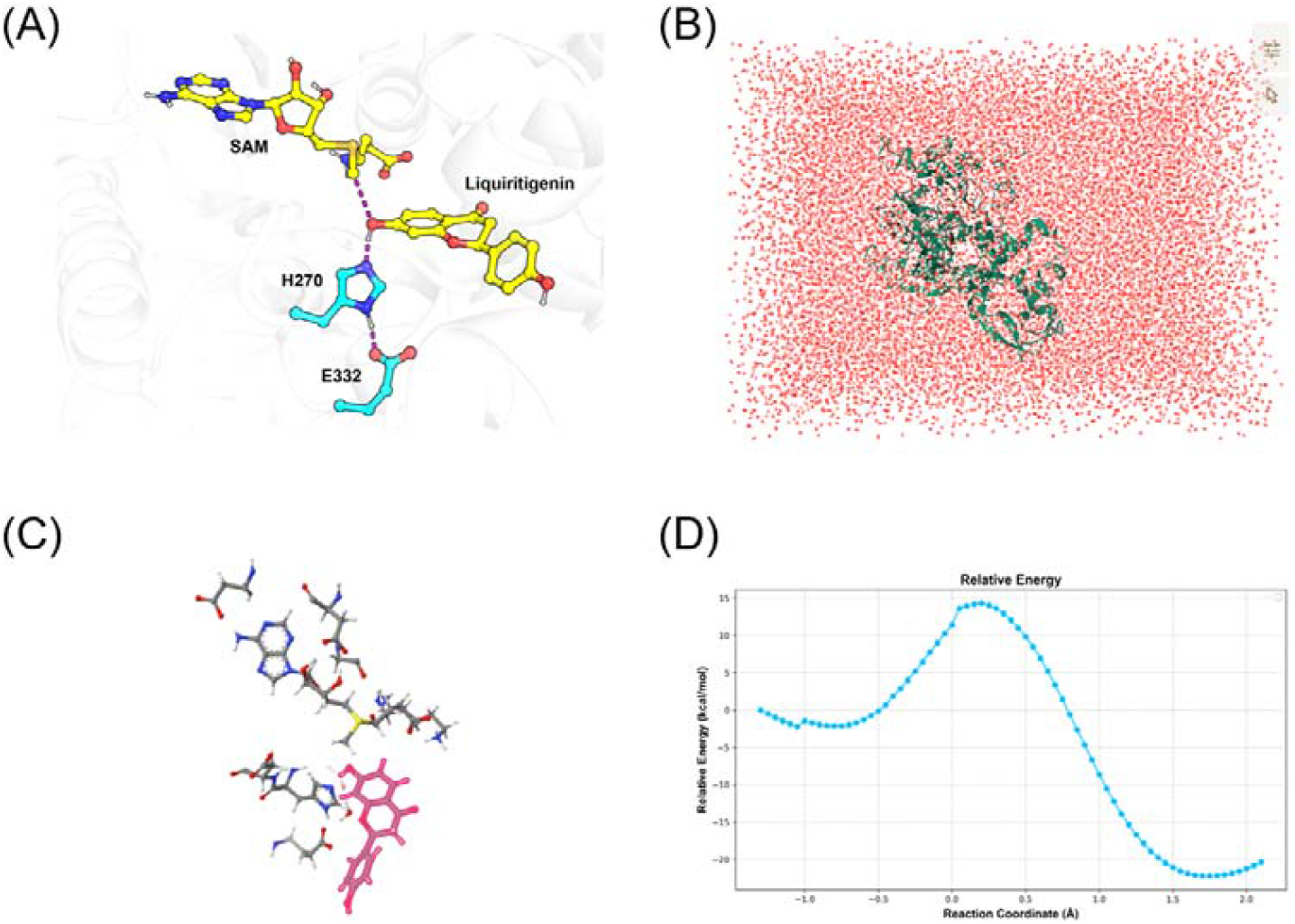
Key computational results generated by EnzySeek for the Isoflavone-4’-O-methyltransferase system. (A) Illustration of the active site in the Isoflavone-4’-O-methyltransferase system. (B) System solvation and force field parameterization performed by EnzySeek. (C) Autonomous partitioning of the QM region by EnzySeek based on manually specified residue indices. (D) Potential energy surface (PES) of the reaction pathway obtained via scanning by EnzySeek.

Initial MD simulations of the system revealed that neither 2,7,4^′^-trihydroxyisoflavanone nor liquiritigenin could maintain suitable reaction distances and orientations with H270. We hypothesized that molecular mechanics force fields were inadequate for describing these specific residue interactions accurately. Consequently, we utilized the EnzySeek Agent to assist in conducting QM/MM MD simulations at the GFN2-xTB level.

1. We provided manually selected frames from the MD trajectories (in PDB format) as input files (Figure 4B).
2. With the assistance of the Agent, we defined the QM region by specifying the corresponding residue numbers (Figure 4C).
3. The Agent automatically configured the input files and launched 16 distinct 100 ps QM/MM MD trajectories starting from different substrate poses.
4. Ultimately, the system was found to remain stable throughout the QM/MM MD trajectories. Subsequent PES scans on key degrees of freedom proved the overall feasibility of the reaction (Figure 4D).

## Discussion

This paper introduces our accelerated framework for enzyme catalysis computations, utilizing an AI Agent to assist human researchers in task execution and result analysis. The results demonstrate that human-Agent collaboration can significantly enhance the efficiency of enzyme catalysis simulations. It drastically reduces the unnecessary cognitive burden on human researchers when submitting tasks and processing results, enabling them to focus their attention on critical scientific questions.

In its current development stage, the EnzySeek Agent is primarily utilized to replace the manual operation of various tools and the subsequent analysis of results. We argue that for experienced senior researchers, invoking tools, setting parameters, and processing outputs are highly repetitive processes; delegating these mechanical operations to an Agent can significantly boost overall efficiency. For beginners just entering the field, integrating different domain-specific tools and deeply understanding the underlying principles of each carries a steep learning curve. Furthermore, the inconsistent operational logic across different software packages is highly prone to causing user errors. Utilizing the Agent can help researchers rapidly navigate the workflow to obtain results, thereby substantially lowering the barrier to entry. For interdisciplinary researchers, who often need to quickly acquire valuable reference data without necessarily delving into the fundamental physics behind every operation, the Agent can bypass complex learning processes, thus promoting cross-disciplinary integration.

EnzySeek is designed to continuously learn expert knowledge and operational habits from novel research literature and historical computational tasks. During operation, EnzySeek evaluates the confidence level of its current decisions. For operations with low confidence or those that are computationally expensive (such as high-throughput tasks or prolonged simulations), the system pauses and awaits human decision-making. As the volume of these human-AI collaborative tasks increases, the system’s decision-making is expected to become increasingly precise, which translates to reduced human intervention throughout the research process. In the future, EnzySeek holds the promise of autonomously completing computational pipelines in the field of enzyme catalysis.

In conclusion, as interest in biosynthesis continues to grow and the enzyme catalysis community flourishes, the field faces increasing computational demands alongside a shortage of specialized personnel. By applying Agent technology to enzyme catalysis simulations, we enable AI to assist humans in task scheduling and result analysis. This paradigm has the potential to substantially elevate the research efficiency of enzyme catalysis simulations, foster the seamless integration of computational and experimental approaches, and ultimately enhance our fundamental understanding and rational engineering of enzymes.

